# Co-isolation of *Penicillium citrinum* and its cell-switching partner *Meyerozyma guilliermondii* from a geothermal power plant

**DOI:** 10.1101/2024.06.19.599737

**Authors:** Danaé Bregnard, Guillaume Cailleau, Wart van Zonneveld, Simona Regenspurg, Saskia Bindschedler, Pilar Junier

## Abstract

Progresses in geothermal energy and deep drilling technologies have opened a new window into the terrestrial subsurface. This provides direct access to deep geothermal fluids used to produce heat and electricity, creating an opportunity to isolate and characterize novel microbial strains from these extreme habitats. In this study, we report the co-isolation of two fungal strains. *Penicillium citrinum* (strain HEK1) was isolated first and thought to be axenic. However, upon exposure to stress (frost and ethanol), a second strain, corresponding to the dimorphic yeast *Meyerozyma guilliermondii* (strain HEK2), appeared in HEK1 cultures. Strain HEK2 appeared first in the cultures and was followed by the subsequent re-growth of strain HEK1, underscoring their close relationship. Moreover, strain HEK2, able to switch from yeast cells to pseudohyphae when growing alone, did not produce pseudohyphae when in direct contact with strain HEK1. Altogether, our results indicate an intricate interaction between these strains that may allow them to thrive in the deep subsurface. These two fungi represent the first fungal strains isolated from deep geothermal fluids. Their presence within the fluids was confirmed through molecular analysis. The isolation of these strains emphasizes the importance of considering fungi when investigating microbial diversity in subsurface geothermal environments.

**Graphical abstract:** 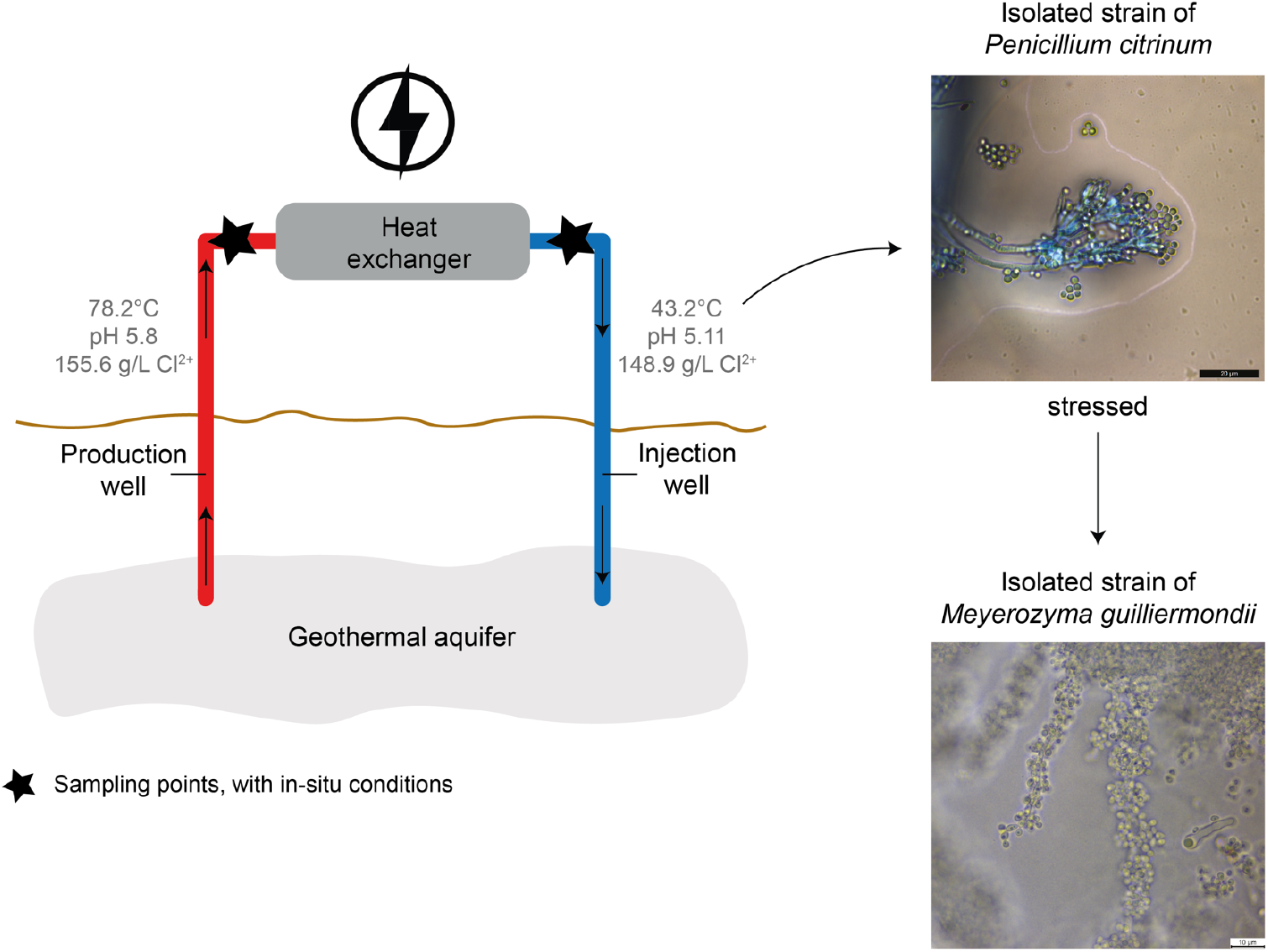

**Highlights:**

- First fungal strains isolated from a geothermal power plant
- The two fungal strains were co-isolated from a geothermal fluid used for heat production
- Surprising isolation of the cell-switching yeast upon stress exposure of an apparently axenic culture of the filamentous fungus
- Fungal strains with high resistance to stressors and no apparent competition for carbon sources

## 1. Introduction

Improvements in sampling technologies and the continuous development of so-called culture-independent molecular methods based on metagenomic or amplicon sequencing, allows nowadays to assess the diversity of life in extreme environments. This includes environments that had not been accessible until recently, such as deep-sea hydrothermal systems or the deep subsurface [1], [2]. Diversity studies in these types of ecosystems highlight the broad range of habitats that sustain life, further expanding our understanding of the environmental parameters restricting the active growth and survival of organisms. However, despite the various advantages provided by culture-independent methods, cultivating strains under laboratory conditions is still an essential complement that allows to further investigate the phenotypic and metabolic traits of extremophilic microorganisms. This provides further insights into the mechanisms allowing the survival and activity of microorganisms under extreme conditions. Moreover, if additional predictions about the metabolic activities and growth patterns of extremophiles can be confirmed through cultivation studies, metabolic prediction models can be adjusted and become more accurate, especially for environments where life exhibits diverse adaptations to extreme conditions, creating a virtuous feedback cycle [3].

Isolating and cultivating new microbial strains also mean potentially isolating microorganisms with yet unknown metabolic activities, for example by producing novel enzymes or enzymes that are active only under specific conditions. The isolation of *Thermus aquaticus* from a geothermal spring is probably one of the most iconic examples so far, as this has led to the development of the famous Taq polymerase still used nowadays in PCR [4], [5]. Hence, those microbial strains therefore hold the potential to be or to produce useful molecules for biotechnology purposes, whether used as a whole or for their metabolites. This potential is notably amplified in the case of extremophilic and extremotolerant microorganisms, whose unique metabolisms and enzymes diverge substantially from those of the more familiar non-extremophiles [6], [7], [8]. Moreover, studying microbial strains from extreme environments may provide information on the evolution and the conditions limiting life as some extreme environments can be considered as analogous to Early Earth. These strains and their properties also inform about the potential to find extraterrestrial life, as the conditions prevailing are characteristic of other habitable planets [9], [10], [11], [12], [13], [14], [15]. Therefore, exploring the diversity of strains that can be isolated from extreme environments holds great potential for new discoveries.

One example of these underexplored extreme environments is deep subsurface geothermal reservoirs used either for heating or for electricity production. Microbial strains isolated from these deep subsurface geothermal fluids include diverse bacterial species such as *Thermus thermophilus* TMY [16], *Anoxybacillus geothermalis* [17], [18], and *Rhodothermus bifroesti* [19], and archaeal strains including *Pyrobacurum* [20] or *Archaeoglobus fulgidus* [21]. However, to the best of our knowledge, no fungi had yet been isolated from deep geothermal fluids. Here, we report the co-isolation of two fungal strains from deep geothermal fluids of the Heemskerk geothermal power plant (NL), one filamentous fungus and one dimorphic yeast. While the isolation work initially led to apparently axenic culture of a *Penicillium citrinum* (strain HEK1), exposure to abiotic stress led to the emergence of a second strain belonging to the *Meyerozyma* genus. These two strains thus represent the first fungal strains isolated from deep geothermal fluids. Both strains could then be isolated as pure cultures which allowed to characterize their metabolic and phenotypic traits alone and combined, both through direct and indirect (via volatiles and exudates) confrontations. With this, the type of interaction that is established between these two strains could be described into more details. Moreover, in parallel to this culture-dependent approach, a culture-independent assessment of the fungal diversity present within this same power plant was done through molecular methods to confirm the presence of both strains within the power plant itself.

## 2. Methods

### 2.1 Sample collection, enrichment culture and strain isolation

Deep geothermal fluids were sampled before and after the heat-exchanger of the geothermal power plant in Heemskerk (NL), within the surface system. The fluids sampled before the heat-exchanger come from the production well and are herein labelled as “production well”. The fluids sampled after the heat-exchanger are re-injected through the injection well and are herein labelled as “injection well”. At the time of the sampling, fluids at the production well had a temperature of 78.2°C at the well-head and a pH of 5.8. At the injection well, the fluids’ temperature was 43.2°C and the pH 5.11. Currently, the fluid temperature at the production well reaches 100-102°C. Moreover, the salinity (expressed as Cl^2-^ concentration) at the production well was 155.6 g/L while it was 148.9 g/L at the injection well [22]. Prior to sampling, sampling and cooling devices, used to cool down fluids to manipulable temperatures, were flushed with geothermal fluids to avoid contamination. For each sampling point, 1 L of fluid was taken in sterile bottles and conserved at 4°C until use. To selectively enrich for fungi, the Emerson YpSs Broth, ½ strength was used (ATCC® Medium 2370 (ATCC®, USA) [23]) supplemented with 50 mg/L of ampicillin [24]. This medium was used in the past to isolate thermophilic fungi from corn grains [25]. Enrichment cultures were composed of 25 mL of culture media inoculated with 1 mL of geothermal fluid and incubated at 22°C in the dark without agitation. After signs of fungal growth, 1 ml of the liquid culture was plated on Emerson YpSs Agar, ½ strength with ampicillin (50 mg/L) and on Malt Agar (MA; malt: 12 g/L, SIOS Homebrewing, CH, agar: 15 g/L, Biolife Italiana, IT). Strains were further purified on MA plates by consecutive culturing. The first filamentous fungal strain isolated from Heemskerk was designated as strain HEK1 while the yeast strain subsequently isolated was designated as strain HEK2.

### 2.2 DNA extractions, PCR, Sanger sequencing and identification

DNA extractions were done using the FastDNA®SPIN kit for Soil (MP Biomedicals, USA). Fungal biomass (vegetative cells and asexual spores if present) were collected from solid cultures using a lanceolate lance and put directly into the Lysis Matrix E Tube of the DNA extraction kit. Quantification of DNA was done using a QuBit™ dsDNA HS Assay Kit with a Qubit 2.0 fluorometer (Invitrogen). For all PCR; each sample consisted of 10.5 µL PCR-grade water, 12.5 µL 2X ALLin™ Red Taq Mastermix (highQu GmbH), 0.2 µM of forward primer, 0.2 µM of reverse primer and 1 µL of 2 ng/µL DNA. Amplifications were done using a Thermo Scientific Arktik thermal cycler. First, a PCR targeting the whole ribosomal internal transcribed spacer (ITS1 and 2, including the 5.8S rRNA) region was done using the Allin™ RedTaq Mastermix, 2x (highQu GmbH) and the primers ITS1F (5’-CTTGGTCATTTAGAGGAAGTAA-3’) [26] and ITS4 (5’-TCCTCCGCTTATTGATATGC-3’) [27], [28]. The amplification cycle was as follow: denaturation (95°C for 1 min); 40 cycles of denaturation (95°C for 15 sec), annealing (59°C for 15 sec) and elongation (72°C for 15 sec); final elongation (72°C for 15 sec). Then, a PCR targeting the small subunit ribosomal RNA region (SSU-rRNA) was done using the Allin™ RedTaq Mastermix, 2x (highQu GmbH) and the primers TW10-F (3’-GCGGTAATTCCAGCTCC-5’) [29] and Nu-SSU-1196R (5’-TCTGGACCTGGTGAGTTTCC-3’) [30]19/06/2024 20:23:00. The amplification cycle was as follow: denaturation (94°C for 2 min); 35 cycles of denaturation (94°C for 10 sec), annealing (56°C for 10 sec) and elongation (72°C for 30 sec); final elongation (72°C for 8 min). Finally, a PCR targeting the large-subunit ribosomal RNA region (LSU-rRNA) was done using the Allin™ RedTaq Mastermix, 2x (highQu GmbH) and the primers LR0R-F (5’-ACCCGCTGAACTTAAGC-3’) [31] and LR6-R (5’-CGCCAGTTCTGCTTACC-3’) [32]. The amplification cycle was as follow: denaturation (95°C for 5 min); 35 cycles of denaturation (94°C for 30 sec), annealing (52°C for 30 sec) and elongation (72°C for 1 min); final elongation (72°C for 15 sec). The amplified PCR products were purified using a MultiScreen® Filter Plates PCR µ96 (Millipore) and sent for Sanger sequencing (Fasteris, Geneva). The obtained forward and reverse sequences were aligned, if possible, using BioEdit Sequence Alignment Editor version 7.2.5 or the alignment tool provided by NCBI before being compared against NCBI BLAST for identification (https://blast.ncbi.nlm.nih.gov/Blast.cgi).

### 2.3 Maintenance of the fungal strains

Both strains were routinely kept at 4°C on MA slants. For the strain HEK1, routine growth was done at 30°C on MA plates using an agar plug of 0.5 cm of diameter colonized by filamentous fungal biomass as a starting inoculum. For the strain HEK2, routine growth was done at room temperature on MA using yeast biomass and inoculated with a platin loop.

### 2.4 Optical and Cryo-SEM microscopy

Optical microscopy images were taken using a Leica optical microscope (Leica DMR fluorescence microscope and Leica DFC7000T camera). A lactophenol cotton blue staining was done to contrast fungal structures. Further micromorphological characterization of the strain HEK1 was done via Cryo-Scanning Electron Microscopy (Cryo-SEM) using a Quanta^™^ FEG 250 microscope. Samples were prepared using a PP3010 Cryo-SEM/Cryo-FIB/SEM Preparation System (QuorumTech, UK) directly from an agar plug of a solid culture containing fungal biomass grown for 7 days.

### 2.5 Stress tolerance of *P. citrinum* HEK1 and isolation of a yeast strain (HEK2)

Asexual spores of the strain HEK1 were collected from a fully colonized plate (10 cm diameter) with 15 mL of Tween® 80 (0.016%) (Merck, DE). The resulting suspension was centrifugated (Sigma 2-16PK) at 3000x*g* for 10 minutes and the supernatant was removed. To rinse the spores thoroughly from any remaining Tween® 80, 5 mL of Milli-Q® Water (Merck Milipore, DE) were added, and the suspension was centrifugated at 3000x*g* for 10 minutes before the supernatant was removed. After this step, spores were re-suspended in 5 mL of sterile Milli-Q® Water. This spore suspension was then exposed to different abiotic stresses: −80°C, - 20°C, UVC exposure (wavelength: 254 nm), Tween® 80 (0.016%), ethanol 70% (EtOH) (Sigma-Aldrich®, DE), UV and ethanol 70% (EtOH) as well as an autoclave cycle (one spore suspension for each stress, if not mentioned otherwise). For −80°C and −20°C, spore suspensions (2 mL, in 2 mL Eppendorf tubes (Eppendorf SE®, DE) were directly exposed to either of the two temperatures for two weeks. After this, 200 µL of exposed spore suspensions were plated on MA in triplicate and incubated at room temperature. For UVC and EtOH stresses, spore suspensions (200 µL) were plated on MA and then exposed to UVC or EtOH after which they were incubated directly. For EtOH exposure, plates were sprayed with EtOH directly after inoculation and air dried for 5 minutes. For the autoclave stress, spore suspensions (10 mL in glass vials) were exposed to a standard autoclaving round (121°C, 20min). As a control, a fraction of the spore suspension (2 mL, in 2 mL Eppendorf tubes (Eppendorf SE®, DE) was re-suspended in physiological water instead of Milli-Q® Water and stored at 4°C for one week, before being plated on MA (200 µl per plate). After stress exposure, all plates were incubated at room temperature to observe growth. This whole procedure was done twice (round 1 and round 2), except for the Tween® 80 (0.016%) stress, which was done only in round 1. Finally, the yeast that grew on plates initially inoculated with spores from the strain HEK1 exposed to cold stresses was isolated and cultivated on MA plates at room temperature and was further designated as the strain HEK2.

### 2.6 Exposure of *M. guilliermondii* HEK2 to stresses: UVC and 70% ethanol

The yeast strain HEK2 was densely plated on MA and exposed either for 15 minutes to UVC, to 70% ethanol or to 15 minutes of UVC combined with 70% ethanol as previously done for the strain P. *citrinum* HEK1 (see section: “2.5 Stress tolerance of *P. citrinum* HEK1”). Growth was observed both macroscopically and microscopically.

### 2.7 Assessment of fungal diversity in the Heemskerk power plant through molecular methods

In parallel to the enrichment cultures, the fungal diversity present in the geothermal fluids of the Heemskerk power plant was assessed. Geothermal fluids from the production and injection wells were sampled in sterile containers (40L per well), filtered through 0.22 µm nitrocellulose membranes (Merck, Ref. GSWP04700) and resulted in a total of 14 filters for the production well and 20 filters for the injection well. More filters were needed to filter 40L of fluids from the injection well due to the rapid clogging of the filters. Each filter was split into two and DNA extracted from each half. Each half filter was either subjected to a spore-enrichment treatment or to the DNA extraction directly, as mentioned in [33]. However, as the goal of this paper was not to differentiate DNA coming from resistant structures such as spores from DNA coming from less resistant structures, the results coming from both methods were considered equally. DNA extractions were done in triplicates with the FastDNA®SPIN kit for soil (MP Biomedicals, USA) with two additional bead-beating steps [33], [34], [35]. Illumina MiSeq sequencing (2 × 300 bp) was done on DNA extracts using the ITS3_KYO2 (GATGAAGAACGYAGYRAA) [36] and ITS4 (TCCTCCGCTTATTGATATGC) [27] primers. Paired-end sequences were processed through QIIME2 [37] and the DADA2 pipeline for denoising, amplicon sequence variants (ASVs) attribution [38], followed by taxonomic annotation using the UNITE database [39] before being treated in RStudio (software version 4.3.1) [40] with the phyloseq package [41]. Relative abundances of the different fungal genera were used for these analyses. The prevalence (presence or absence in the half-filters) of the genera of interest were calculated (number of half-filters where the genus is present/ number of half-filters analysed).

### 2.8 Confrontations of *P. citrinum* HEK1 and *M. guilliermondii* HEK2

Direct confrontations of the fungal strains HEK1 and HEK2 were done on MA, at room temperature. An agar plug containing biomass of HEK1 was inoculated on one side of the plate (10 cm diameter) while a line (approximately 6 cm in length) of biomass of HEK2 was inoculated in the middle of the other half of the plate. Moreover, to assess the impact of the volatiles produced by one fungal strain on the other and reciprocally, 2-chambers 10 cm diameter Petri dishes (Greiner Bio-One, AT) with malt-agar (MA) medium were used. To assess the impact of exudates produced by the strain *P. citrinum* HEK1 on the growth of the yeast strain *M. guilliermondii* HEK2, liquid cultures of *P. citrinum* HEK1 was grown in malt-broth (MB, malt: 12 g/L, SIOS Homebrewing, CH) at room temperature for 7 days. The resulting culture was then filtered through a 0.22 μm nitrocellulose membrane to keep only the spent medium containing the exudates. The spent medium was mixed with sterile water-agar medium (WA) (before solidification point, approximate temperature of 40°C) at a proportion of 1:1 (final agar concentration: 1.5%) before being poured into 3 cm diameter Petri dishes. Moreover, the same medium (WA) was supplemented with malt (12 g/L, SIOS Homebrewing, CH, agar: 15 g/L, Biolife Italiana, IT), resulting in a water-agar-malt medium with exudates.

### 2.9 Dimorphic growth assessment of the strain HEK2 using agar pads, solid media and volatiles and exudates exposure

Time lapse growth of the yeast cells of HEK2 was recorded using a reversed optical microscope (EVOS M5000, Invitrogen by Thermo Fisher Scientific Inc., USA) taking an image every minute for 96 h on agarose pads made of malt and low-melting point agar (malt; 12 g/L, SIOS Homebrewing, CH; agarose, 10 g/L, TopVision Low Melting Point Agarose, Thermo Fisher Scientific Inc., USA) as described previously [42]. A yeast cells suspension was prepared in MB and 2 µl of this suspension were inoculated in the center of the agarose pad. Furthermore, the expression of yeast cells or pseudohyphal growth were assessed on different solid media that were selected in order to be as limiting as possible for fungal growth: N-free medium (Supplementary Material: Media Composition), *Aspergillus* Minimal Medium (AMM) supplemented with sodium chloride, acetate and propionate (Supplementary Material: Media Composition), water-agar medium (WA) (agar, 15 g/L, Biolife Italiana, IT).

### 2.10 Characterization of carbon use by *P. citrinum* HEK1 and *M. guilliermondii* HEK2

In order to assess the ability to use different carbon (C) sources in either of the two isolated strains, Biolog EcoPlate^™^ (Biolog) containing 31 different carbon sources in triplicates were inoculated with asexual spore suspension for HEK1 (2.1*10^5^ spores/mL) and yeast cells suspension for HEK2 (3.1*10^7^ cells/mL). Effective growth on either of the 31 C sources was spectrophotometrically assessed by measuring the reduction of a tetrazolium dye (included in the plate) at an absorbance of 590 nm using an Agilent BioTek Cytation^™^ 5 Imaging reader. Measures were done at T0 (i.e. directly after inoculation) and after 24, 48, 72, 96 and 120 h of incubation at room temperature. For both organisms, all absorbance values above 0.1 were considered as positive (after T0 absorbance removal). Growth curves were plotted in the RStudio (software version 4.3.1) [40] using the *gcplyr* package [43].

## 3. Results

### 3.1 Isolation of the strain HEK1

A fungal strain (HEK1) was isolated at 22°C from enrichment cultures carried out with geothermal fluids sampled from the injection well of the Heemskerk power plant (fluid temperature: 43.2°C). The fungal strain produced a large amount of white mycelium and green spores on MA plates both at room temperature and at 30°C. At 30°C, the production of yellow pigments was also observed (Figure 1A). Optical microscopy and Cryo-SEM microscopy of the isolate allowed its identification as a fungus belonging to the *Penicillium* genus, due to the presence of characteristic conidiophores made of phialides cells (Figure 1B, C). The strain was further identified as *Penicillium citrinum* through Sanger sequencing. For the ITS rRNA gene region, its closest relatives were *Penicillium griseofulvum* (Query cover: 100%, % identity: 99.59%), *Penicillium citrinum* (Query cover: 100%, % identity: 99.39%) and *Penicillium* sp. (Query cover: 100%, % identity: 99.39%), and *Meyerozyma* sp. (Query cover: 100%, % identity: 100%), with most sequences belonging to *Penicillium citrinum*. For the SSU rRNA region, its first closest relatives were *Penicillium citrinum* (Query cover: 100%, % identity: 99.82%) (most sequences), *Penicillium* sp. (Query cover: 100%, % identity: 99.82%), *Penicillium chrisogenum* (Query cover: 100%, % identity: 99.82%), *Penicillium decumbens* (Query cover: 100%, % identity: 99.82%) and *Penicillium griseofulvum* (Query cover: 100%, % identity: 99.82%). For the LSU rRNA region, the closest relatives were *Penicillium citrinum* (Query cover: 100%, % identity: 99.27%) (most sequences) and *Penicillium* sp. (Query cover: 100%, % identity: 99.14%). Accordingly, the strain HEK2 was considered as belonging to the *Penicillium citrinum* species complex and was thus further designated as *P. citrinum* HEK1.

**Figure 1:**
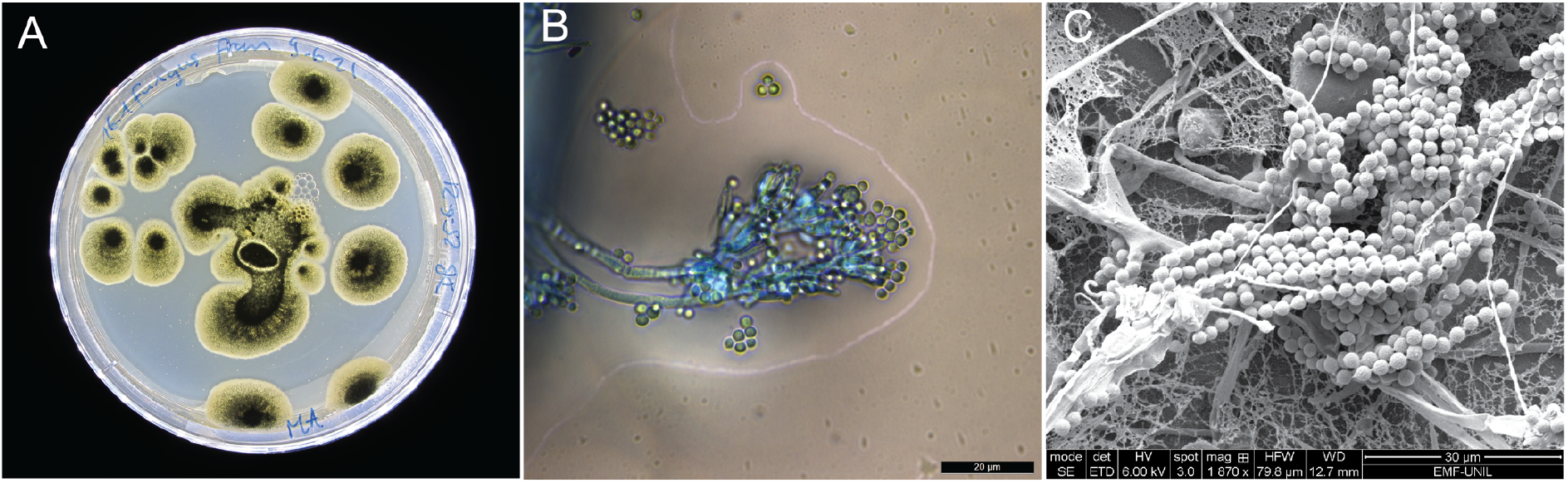
A) *Penicillium citrinum* grown on MA at room temperature. B) Optical microscopy of *Penicillium citrinum* HEK1 grown on MA at 30°C and stained with lactophenol blue. Scale bar represents 20 µm. D) Cryo-SEM images of the fungus *Penicillium citrinum* HEK1 grown on MA at 30°C. Scales bar represent 30 µm.

### 3.2 Stress tolerance of *P. citrinum* HEK1 and isolation of a yeast strain (HEK2)

The resistance of spores of *P. citrinum* HEK1 to various stressors was assessed. Spores, of *P. citrinum* HEK1 were collected and exposed to individual or combined stresses including UV, 70% ethanol, Tween® 80, frost (−20°C and 80°C), and autoclaving (combined pressure and temperature). Spores that were challenged were compared to a control in which spore suspensions were kept at 4°C in a physiological solution and not exposed to any stresses before being plated on MA plates. Growth from stressed spore suspension was assessed after 11 days post inoculation (DPI). Except autoclaving, none of the stresses applied prevented *P. citrinum* HEK1 re-growth (Figure 2A). In addition, when exposed to −20°C, −80°C, 70% ethanol, and Tween® 80 (0.016%) at 4°C, as well as in one replicate of the control, a yeast appeared on the plates alongside the sporulating mycelium of *P. citrinum* HEK1 (round 1) (Figure 2B). The experiment was repeated to confirm these results (round 2), and the growth of yeast cells was once more observed in the −20°C and −80°C and the 70% ethanol treatments (Table 1, Supplementary Figure 1). Moreover, as growth was followed more precisely on plates inoculated with the stressed spore suspensions, it was observed that the initial growth of the yeast was followed by growth of *P. citrinum* HEK1 and not the other way around (Supplementary Figure 1). Exposure of HEK2 to UVC and 70% ethanol, either separately or combined, did not prevent the growth of the yeast (Supplementary Figure 2), but the colonies changed in texture (macroscopical smoothness of the colony appeared visually different) after exposure to UVC or to UVC combined with ethanol. Microscopical images showed no difference between the unstressed yeast cells, the cells exposed to UV and ethanol, and the cells exposed to ethanol only (Supplementary Figure 3). Moreover, cells that had been exposed to UV and ethanol displayed again a regular macroscopical colony smoothness once re-plated on a fresh MA plate.

**Table 1:** Table summaryzing the results obtained in the stress exposure experiment of strain HEK1 (rounds 1 and 2), growth of strain HEK1 is indicated along to the appearance of yeast cells upon each stress treatment.

| Stress | Round 1 |  | Round 2 |  |
| --- | --- | --- | --- | --- |
|  | HEK1 | Yeast cells | HEK1 | Yeast cells |
| -20°C | Yes | Yes | Yes | Yes |
| -80°C | Yes | Yes | Yes | Yes |
| UVC | Yes | No | Yes | No |
| EtOH 70% | Yes | Yes | Yes | Yes |
| UVC + EtOH 70% | Yes | No | Yes | No |
| Tween® 80 (0.016%) at 4°C | Yes | Yes | NA | NA |
| Autoclaving cycle | No | No | No | No |
| No stress | Yes | No (2 replicates);<br>Yes (1 replicate) | Yes | No |

**Figure 2:**
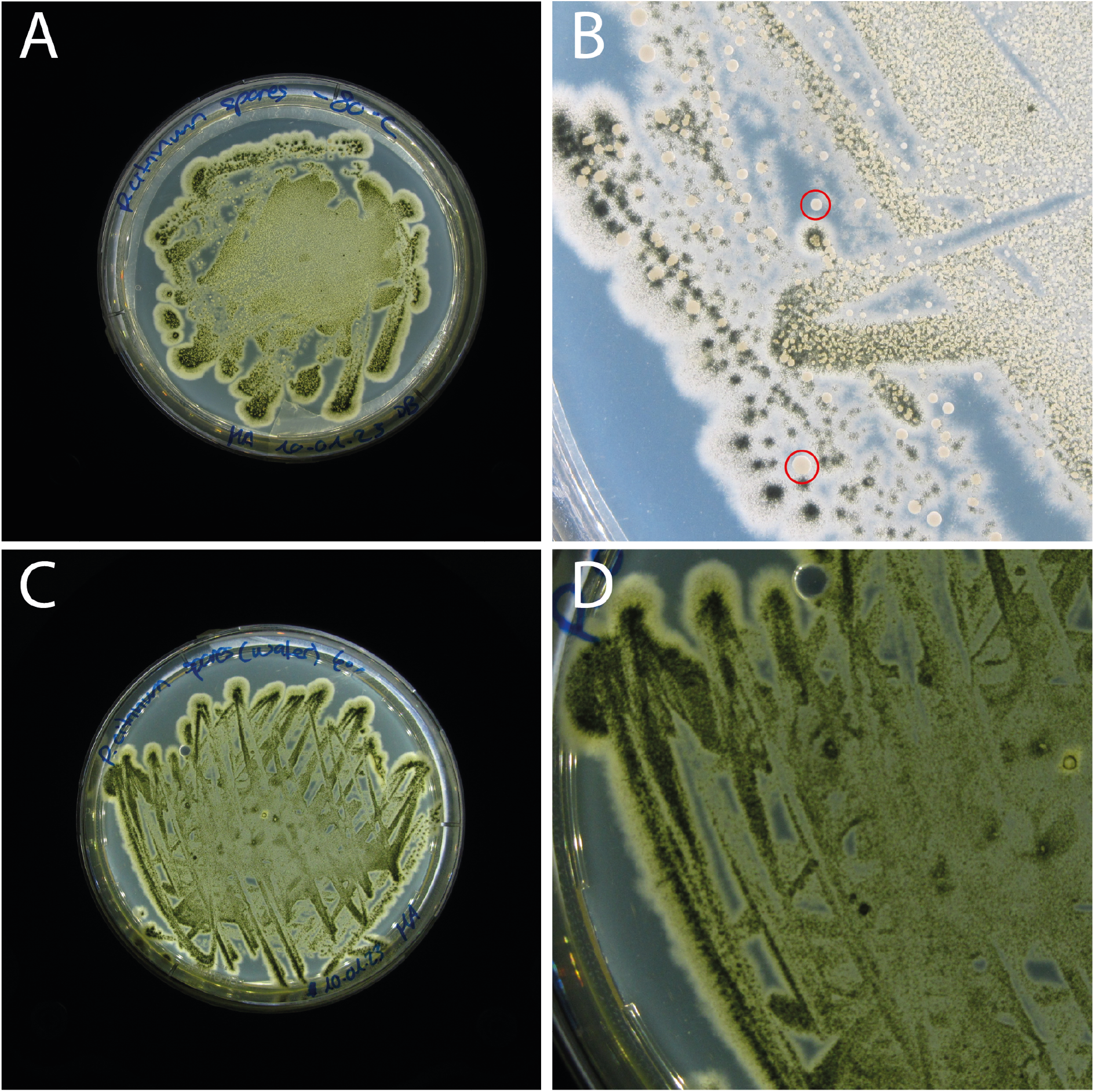
Comparison of growth upon stress after 14 days-post inoculation. A) Culture coming from spores of *P. citrinum* HEK1 in MilliQ water after exposure to −80°C. B) Magnification of the culture in A, where both *P. citrinum* HEK1 and the yeast strain HEK2 (red circles) can be observed. C) Culture from the same initial spore suspension used in A but resuspended in physiological water instead of MilliQ water as control. D) Magnification of the culture in C, where only *P. citrinum* HEK1 (hyphae and spores) can be observed.

**Figure 3:**
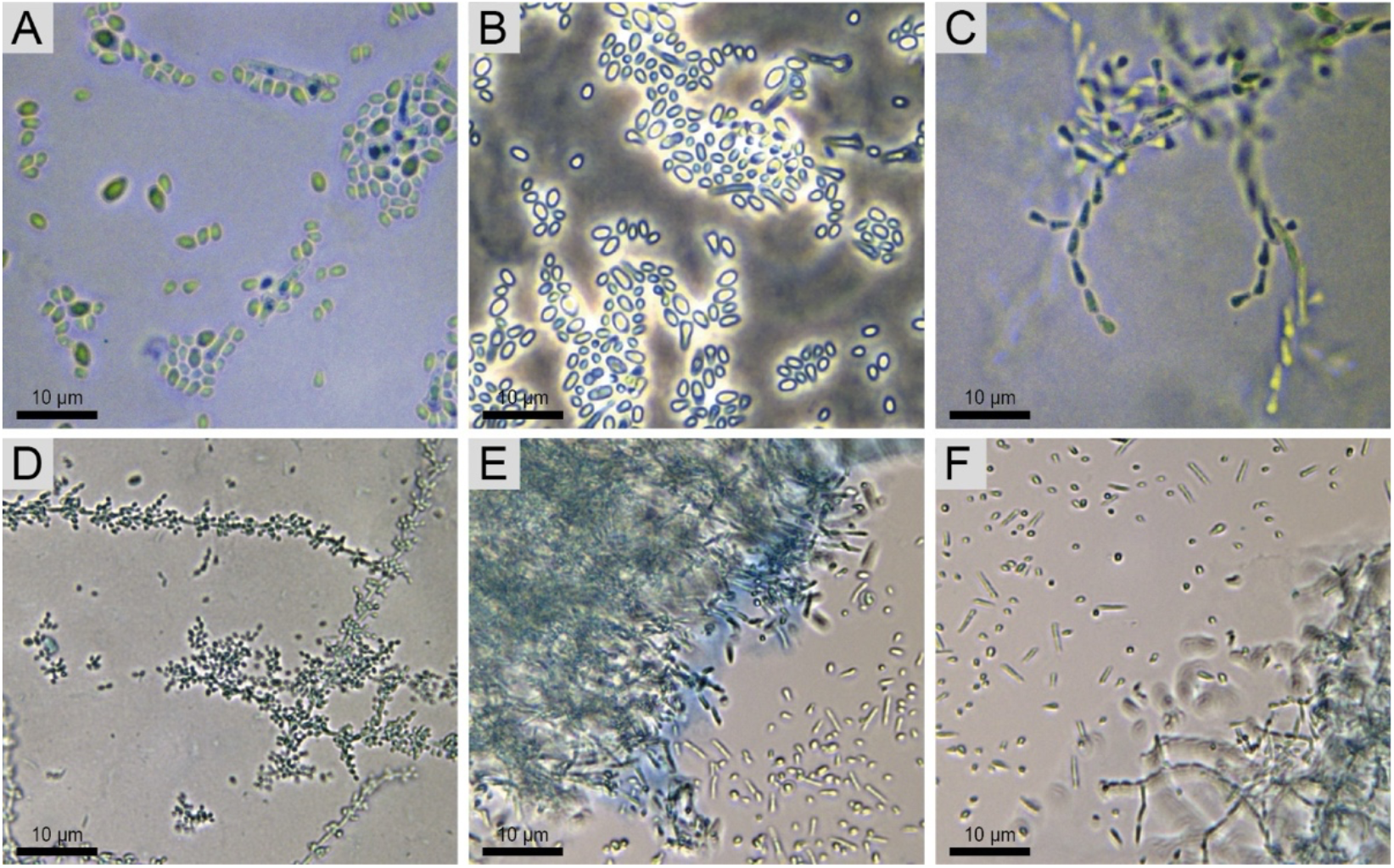
Optical microscopy images of the different cell types of M. guilliermondii HEK2. A) Yeast cells B) Yeast cells under phase contrast C) Pseudohyphae D) Budding pseudohyphae E-F) Interphase between yeast cells and pseudohyphae. Scale bars represent 10 μm.

### 3.3 Identification of the yeast strain HEK2 and co-existence with strain HEK1

The yeast isolated from stressed spore suspensions of strain HEK1 was identified as *Meyerozyma* sp. through Sanger sequencing. For the ITS rRNA gene region, its closest relatives were *Meyerozyma guilliermondii* (Query cover: 100%, % identity: 100%), an uncultured *Meyerozyma* (Query cover: 100%, % identity: 100%), and *Meyerozyma* sp. (Query cover: 100%, % identity: 100%), with most sequences belonging to *Meyerozyma guilliermondii*. For the SSU rRNA region, its first closest relatives were *Meyerozyma guilliermondii* (Query cover: 100%, % identity: 99.6%), *Meyerozyma caribbica* (Query cover: 100%, % identity: 99.6%) (most sequences), *Meyerozyma carpophila* (Query cover: 100%, % identity: 99.6%), *Meyerozyma smithsonii* (Query cover: 100%, % identity: 99.6%) and an uncultured fungus (Query cover: 100%, % identity: 99.6%). For the LSU rRNA region, only the sequence obtained with the LR0R primer was used, as the sequence obtained with the LR6f primer was of very low quality. The closest relatives to the LR0R sequence were *Meyerozyma guilliermondii* (Query cover: 100%, % identity: 100%) (most sequences) and one sequence of *Meyerozyma caribbica* (Query cover: 100%, % identity: 100%). Accordingly, the strain HEK2 was considered as belonging to *Meyerozyma guilliermondii* (anamorph – basionym: *Pichia guilliermondii* Wick) reference needed. Macroscopic and microscopic observations confirmed that the strain HEK2 formed yeast cells and pseudo-hyphae (Figure 4).

**Figure 4:**
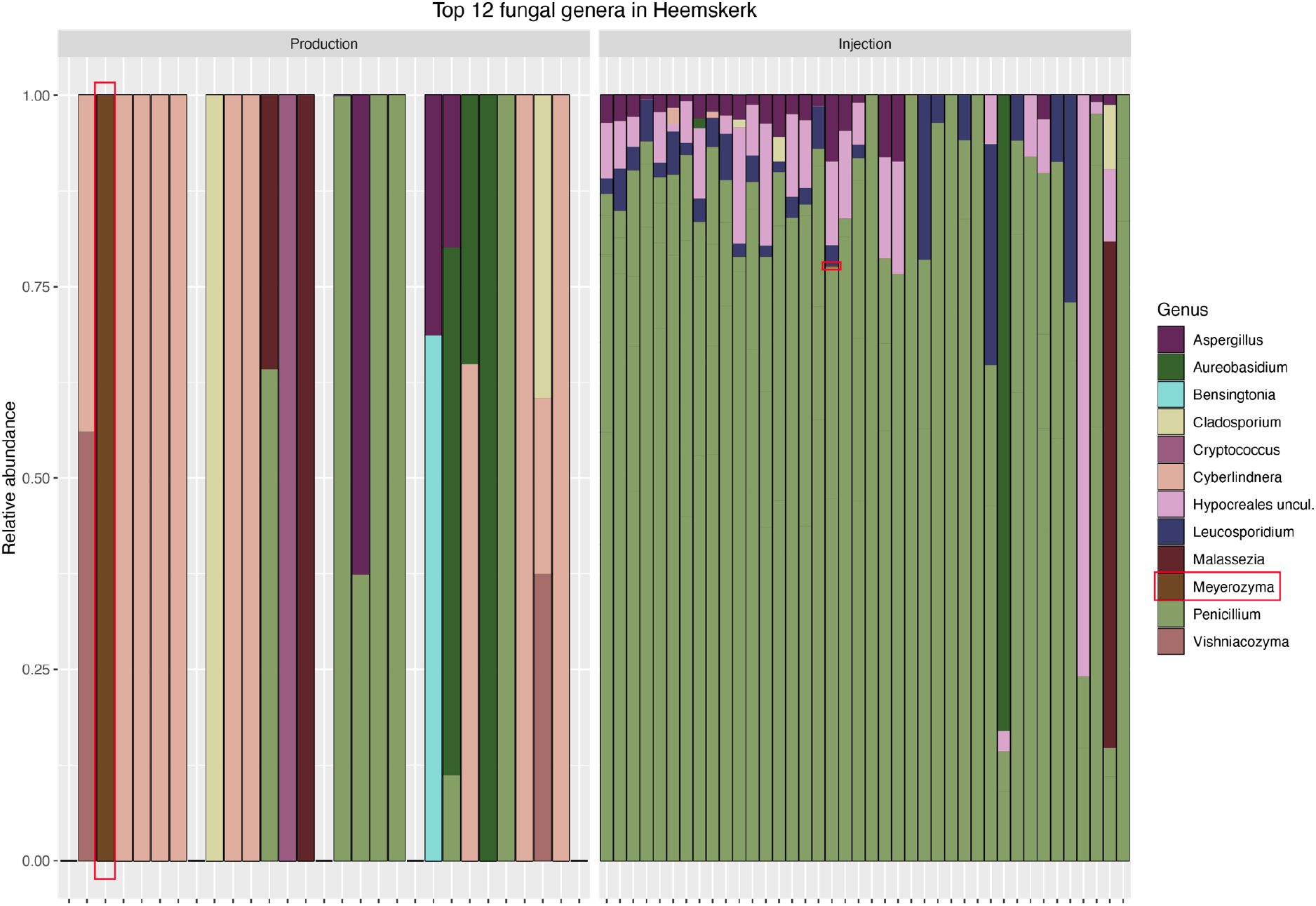
Top 12 fungal ASVs detected in the Heemskerk geothermal power plant through molecular methods at the production (before the heat-exchanger) and injection (after the heat-exchanger) wells. The *Meyerozyma* genus is highlighted in red. The *Penicillium* genus can be seen in light green.

### 3.4 Presence of the *Penicillium* and *Meyerozyma* genera in the fluids detected through molecular analysis

*Penicillium* was present in 25% (7/28 half-filters) of the samples at the production well and it was the dominant genus in 100% of the filters analyzed from the injection well (present in all the 40 half-filters) (Figure 4). ITS sequences affiliated to *Meyerozyma* were also present at the production well and at the injection well, but only in one half-filter in each well. In the sample in which it was detected in the production well, *Meyerozyma* corresponded to the near totality of the genera identified (relative abundance).

Confrontations of the two organisms in MA showed that neither *P. citrinum* HEK1 nor *M. guilliermondii* HEK2 displayed any sign of stress or of growth impairment when co-cultured and the strain HEK2 consistently grew as a yeast (Figure 5). In addition, closer analysis of the Cryo-SEM images revealed unknown round structures (approximate size: 10 – 25 µm) (Figure 5D), that could be compatible with the size of sexual reproductive structures of the yeast and could correspond to *M. guilliermondii* HEK2, but this cannot be confirmed directly.

**Figure 5:**
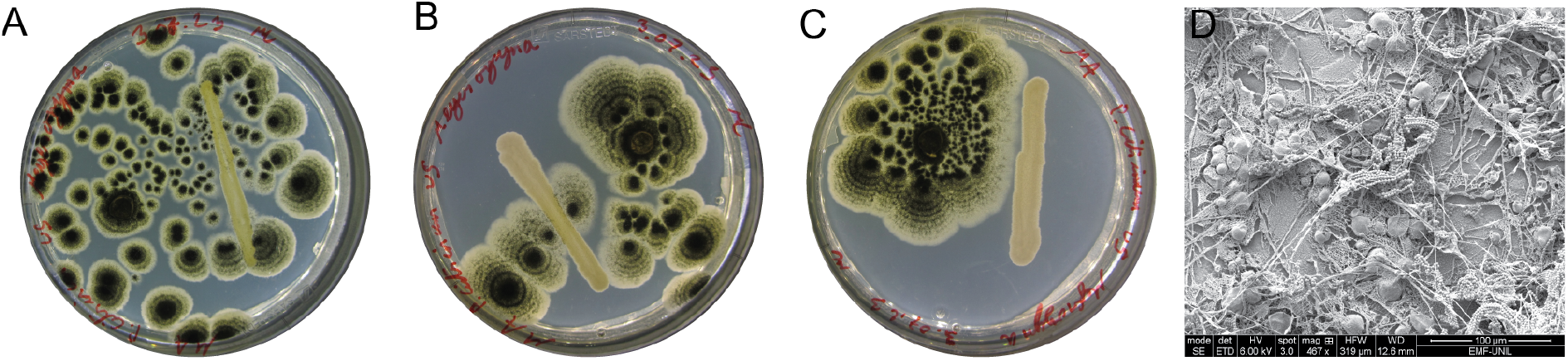
Direct confrontations between *P. citrinum* HEK1 and *M. guilliermondii* HEK2 on MA plates in triplicates (A to C) after 11 DPI. D) Cryo-SEM images of the fungus *Penicillium citrinum* HEK1 grown on MA at 30°C with potential yeast structures. Scales bar represent 100 µm.

### 3.5 Morphological switch of the yeast strain HEK2

During the initial isolation and in the direct confrontation assays with HEK1, *M. guilliermondii* HEK2 was only observed to form yeast cells. However, it always produced pseudohyphae when cultivated alone on solid MA medium (Figure 6A-B). Pseudohyphal formation was confirmed through microscopy (Figure 6B) and corresponded to macroscopical features observed at the border of colonies (Figure 6A). Growth on agar pads (MA) was monitored during 96 h using an inverted microscope. Yeast cells were observed to elongate and produce pseudohyphae only at the border of the colony (Figure 6C-D). Moreover, the formation of septa within the pseudohyphae was recorded (Supplementary Figure 5).

**Figure 6:**
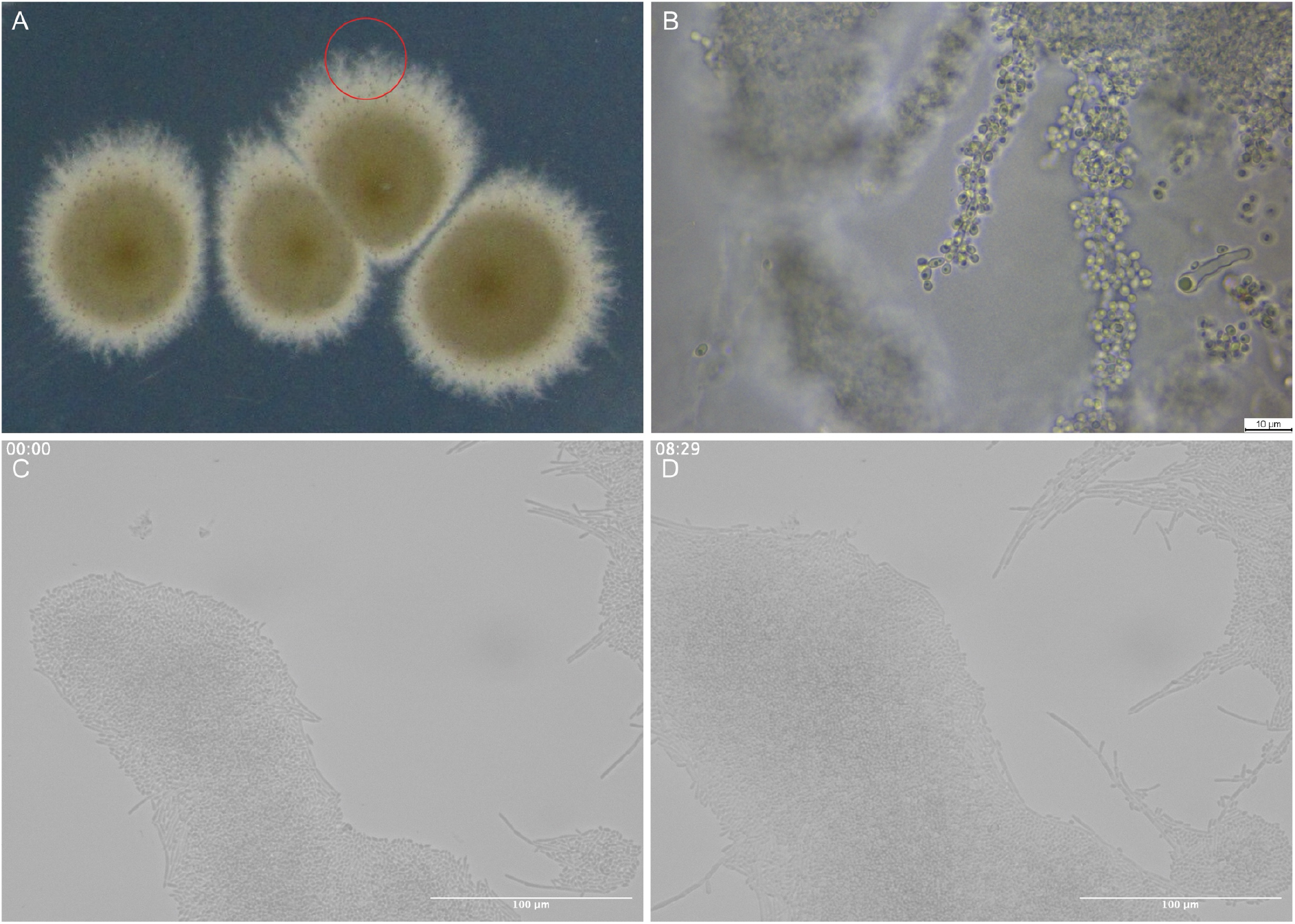
Macroscopic (A) and microscopic (B-D) images of *M. guilliermondii* HEK2 pseudohyphal growth on MA and on an agarose pad containing malt. A) Pseudohyphae can be observed at the border of the colony (red circle). B) Microscopical image of the pseudohyphae located within the red circle of the image A. Scale bar represents 10 µm. C-D) Time lapse images of the growth of yeast cells into pseudohyphae of *M. guilliermondii* HEK2 on an agarose pad containing malt at two time points: 24h (C) and 32h29 (D) post-inoculation. Scale bar represents 100 μm.

### 3.6 Effect of different media on the dimorphic growth of HEK2

To further investigate the factors triggering the formation of pseudohyphae, yeast cells were inoculated on different solid media (N-Free, supplemented AMM and water-agar (WA) (Figure 7A-C). The strain HEK2 was able to grow on the three media, albeit with a visually slower growth than on MA plates. Furthermore, the strain produced few pseudohyphae on N-Free medium, many pseudohyphae on supplemented AMM medium, and small colonies with no pseudohyphae on WA.

**Figure 7:**
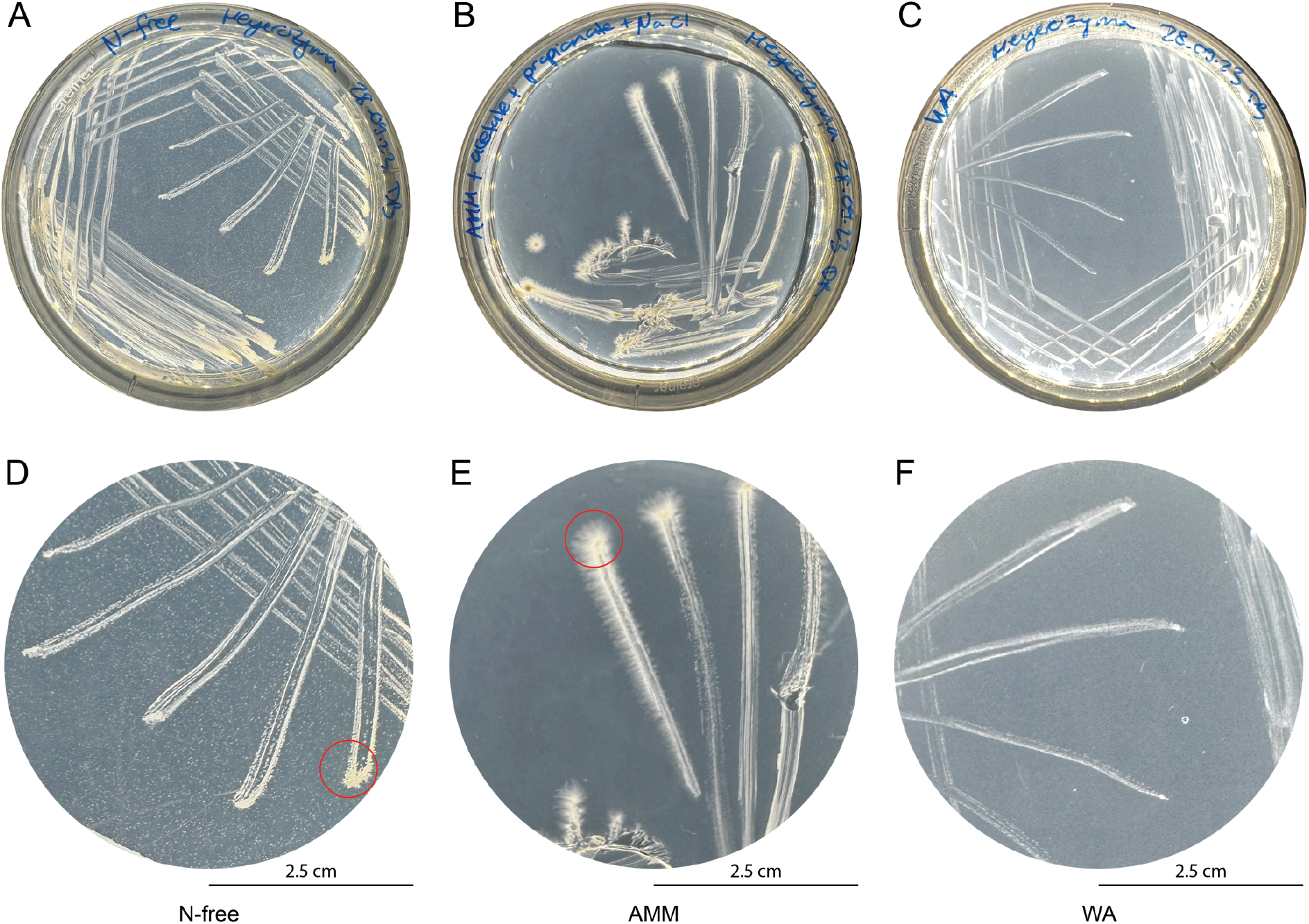
Macroscopic images of *M. guilliermondii* HEK2 on different solid media. A-C) Full plates. D-F) Magnification of the plates. Red circles highlight pseudohyphal formation. A) Growth on N-free medium with the presence of yeast cells and pseudohyphae. B) Growth on supplemented AMM medium with the presence of yeast cells and pseudohyphae. C) Water-agar (WA) medium with only yeast cells. D) N-free medium. E) AMM medium F) Water-agar medium (WA).

### 3.7 Effect of HEK1 volatiles or exudates on the growth of HEK2

Afterwards, to test whether volatile or exudate compounds produced by *P. citrinum* HEK1 would influence the growth of *M. guilliermondii* HEK2, both organisms were plated on split MA plates, limiting physical contact. This showed that in the absence of contact, *M. guilliermondii* HEK2 grew and produced pseudohyphae (Figure 8A). On plates (WA and WAM) prepared with filtered exudates produced by *P. citrinum* HEK1, *M. guilliermondii* HEK2 produced pseudohyphae (WAM (Figure 8B) and WA (Figure 8C)). The density of pseudohyphae could only be assessed macroscopically (Figure 7) but not quantified microscopically, as microscopical slides preparation destroys the density of pseudohyphae.

**Figure 8:**
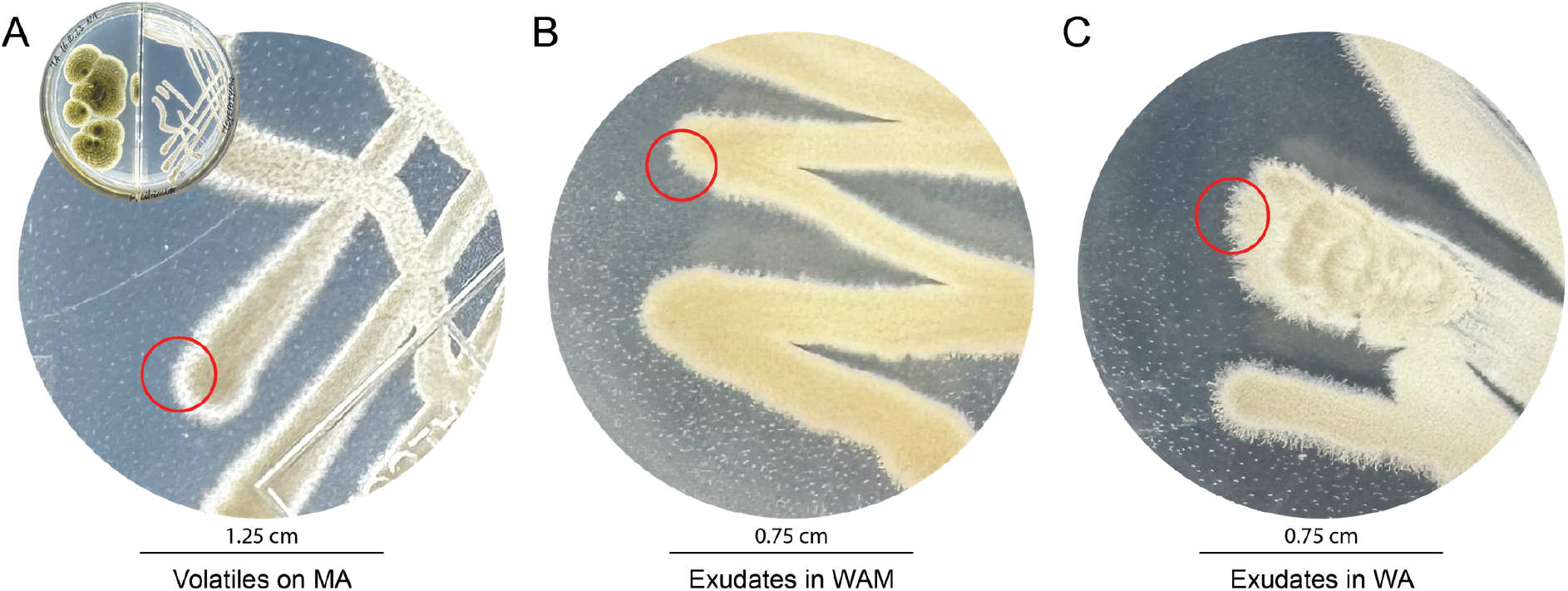
Macroscopic images of *M. guilliermondii* HEK2 during exposure to volatiles or exudates of *P. citrinum* HEK1. Red circles highlight pseudohyphal formation. A) Volatiles experiment between *P. citrinum* HEK1 (left) and *M. guilliermondii* HEK2 (right) on MA (malt-agar) in split Petri dishes at 15DPI. B-C) Exudates experiment between *P. citrinum* HEK1 and *M. guilliermondii* HEK2 15 DPI on WAM: water-agar-malt (B) and on WA: water-agar (C).

### 3.8 Characterization of carbon use by *P. citrinum* HEK1 and *M. guilliermondii* HEK2

To characterize carbon usage patterns, Biolog EcoPlates^™^, which allowed comparing the use of 31 common carbon sources in both strains, were used. After 120 h of incubation, *P. citrinum* HEK1 was shown to be able to grow using Tween40, Tween80, glycogen, D-cellobiose, β-methyl-D-glucoside, D-xylose, i-erythritol, D-mannitol, N-acetyl-D-glucosamine, D-galacturonic acid, 2-hydroxy benzoic acid, 4-hydroxy benzoic acid, γ-amino butyric acid, D-malic acid, L-arginine, L-asparagine, L-phenylalanine, L-serine, L-threonine, glycyl-L-glutamic acid, phenylethyl-amine and putrescine as sole carbon sources (Table 2, Supplementary Figure 5). *M. guilliermondii* HEK2 was able to grow using a smaller set of carbon sources (pyruvic acid methyl ester, Tween40, Tween80, D-cellobiose, β-methyl-D-glucoside, D-xylose, i-erythritol, D-mannitol, N-acetyl-D-glucosamine, γ-amino butyric acid, D-malic acid, L-arginine, L-asparagine and L-serine). It was also observed to grow in glycogen, but the absorbance values were low (Table 2, Supplementary Figure 5). Considering only the carbon sources that clearly sustained growth of the two strains, the strain HEK1 was able to use a wider range of carbon sources (D-galacturonic acid, 2-hydroxy benzoic acid, 4-hydroxy benzoic acid, L-phenylalanine, L-threonine, glycyl-L-glutamic acid, phenylethyl-amine, and putrescine), compared to HEK2, which was able to use only one carbon source (pyruvic acid methyl ester) that seems to be unavailable to *P. citrinum* HEK1.

**Table 2:**
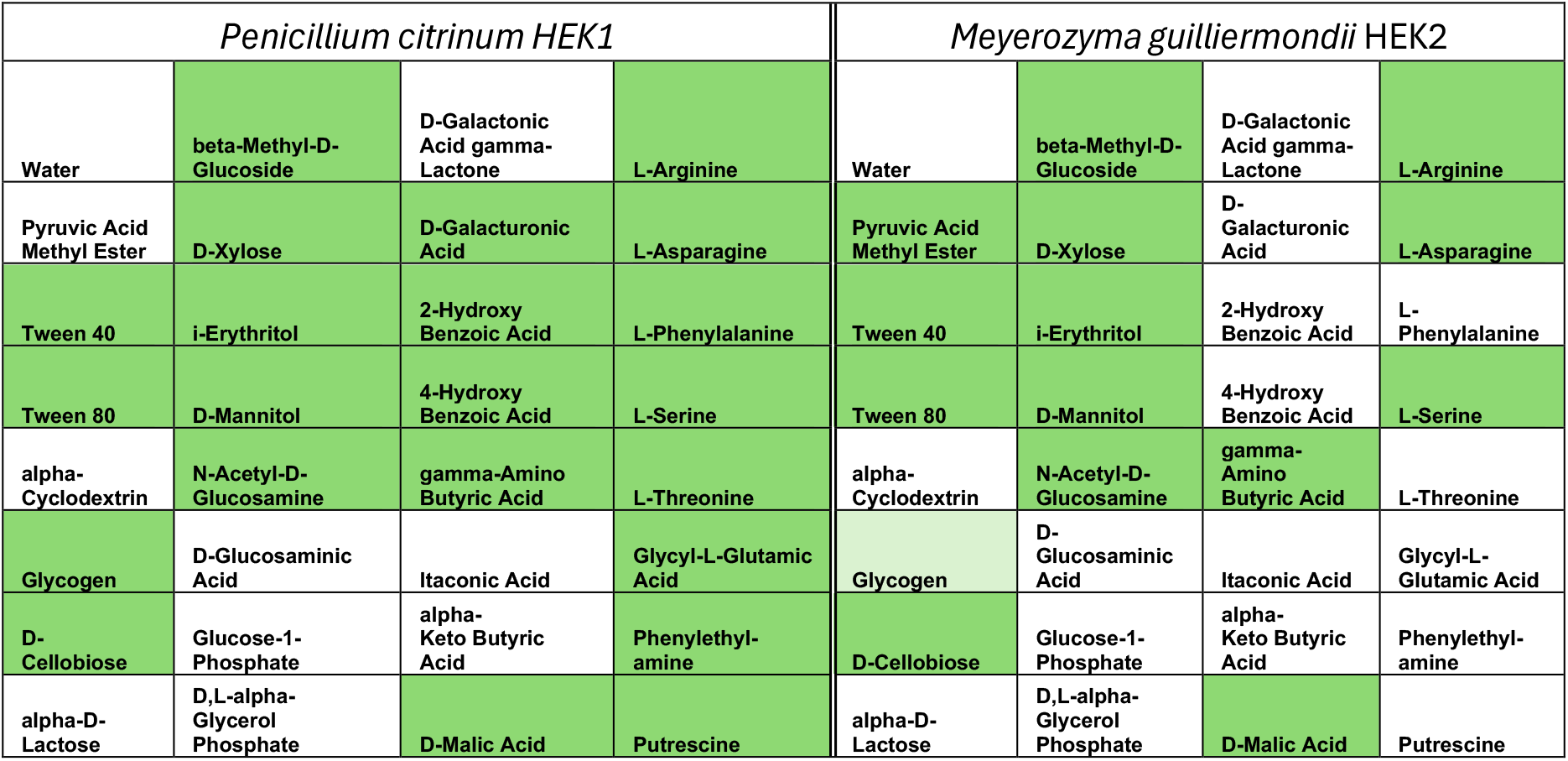
Summary of the carbon (C) sources sustaining the growth of the strain HEK1 and the strain HEK2. Green: high absorbance values and clear visual growth. Light-green: growth observed visually, with low absorbance values.

## 4. Discussion

In this study, two fungal strains were co-isolated from geothermal fluids used for heat production. The first strain was isolated from the surface-system leading to the injection well of the Heemskerk geothermal power plant and belonged to the *Penicillium citrinum* species complex [44]. The *Penicillium* genus, which contains at least 354 different species, is very common in all types of environments and is well-known for its penicillin producing members [45], [46]. The wide distribution of the *Penicillium* genus, including in extreme environments such as deep-sea sediments, salterns, geothermal hot springs, and deep-sea hypersaline basins [47], [48], [49], [50], makes it a good candidate to be found within deep geothermal fluids. The resistance of the *P. citrinum* HEK1 to the various stressors tested (UV, ethanol, cold stresses, and surfactant exposure) highlights its ability to survive to extreme conditions such as those in the geothermal power plant. *P. citrinum* is described as a mesophile and has been used for enzyme production for food processes [51], the production of copper oxide nanoparticles [52], as well as for the production of biomass for the biosorption of different metals, such as lead [53], uranium [54] and copper [55]. This species is known to produce citrinin, a pigmented nephrotic neurotoxin, and tanzawaic acid A, an antifungal compound [56], [57].

The subsequent isolation of a yeast strain from apparently axenic cultures of *P. citrinum* HEK1 cultures exposed to stress is remarkable and to our knowledge is presented here for the first time. The ascomycetous yeast *M. guilliermondii*, formerly called *Pichia guilliermondii*, is a species complex that includes the species *Meyerozyma guilliermondii, Meyerozyma carpophila* and *Meyerozyma caribbica* [58]. *M. guilliermondii* is a saprophytic yeast common in the environment that can be an opportunistic human pathogen [58]. Strains from *M. guilliermondii* have been isolated from a wide variety of environments [59], including deep-sea hydrothermal ridges [60] and dishwashers [61], [62], which are notably known to host extremophilic yeasts [61], [62]. This non-pigmented yeast is known to be tolerant to multiple stresses including salt (16% NaCl [63] and 18% KCl [64]), copper [65] and manganese (5494 ppm) [64]. It has moreover been widely used in biotechnological applications, as it is able to use various substrates for growth and to produce several useful compounds [66]. The wide diversity of environments from which it has previously been isolated as well as its tolerance to several stressors seem to allow the yeast *M. guilliermondii* to survive in geothermal power plants as well.

Furthermore, the capacity of the strain *M. guilliermondii* HEK2 to switch from yeast cells to pseudohyphae provides an additional mechanism of resistance and further enhances its adaptation to changing environmental conditions. Indeed, *M. guilliermondii* is the current name of the opportunistic fungal pathogen known as *Candida guilliermondii* (anamorph name) or *Pichia guilliermondii* (teleomorph name) [67], which is known to switch from yeast cells to pseudohyphae [68]. In *C. guilliermondii*, such phenotypic switching also involves a pigmentation change (from white to brown) and the switch from yeast cells to pseudohyphae has been shown to increase the resistance of the strain to antifungal compounds [68]. Cell morphology switching is reversible and known to be induced in most cases by environmental factors [68]. Its role in pathogenicity is also well known for other *Candida* species. However, pathogenicity is not the only role of phenotypic switching and in *Candida albicans* pseudohyphae are used to enhance nutrient scavenging [69]. Such mechanism is probably relevant for *M. guilliermondii* HEK2 as well, as the switch from yeast cells to pseudohyphae was observed when the yeast cells were not growing, but space and nutrients were still available further away in the culturing plate. Pseudohyphae seemed thus to be formed to allow the strain to propagate further into its environment.

The interaction between the *P. citrinum* HEK1 and *M. guilliermondii* HEK2 was further investigated. Direct confrontations on agar plates revealed that the presence and the contact of the other organism did not impair their reciprocal growth but that *M. guilliermondii* remained as a single-celled yeast. No negative interactions were observed even though these two strains were able to use similar carbon sources. Furthermore, growth in both strains did not change when the two organisms grew in indirect contact through volatiles of the other strain. Conversely, the growth of *M. guilliermondii* HEK2 on water-agar containing exudates of *P. citrinum* HEK1 resulted in the production of a fair amount of biomass and pseudohyphae at the borders of the colonies, while on water-agar alone, *M. guilliermondii* HEK2 was barely able to grow and did not produce any pseudohyphae. Therefore, the exudates produced by the strain *P. citrinum* HEK1 in malt-broth provide sufficient nutrients for *M. guilliermondii* HEK2 to grow and be able to produce pseudohyphae to further colonize the medium. The difference between the nutrients left from the initial medium and the exudates themselves were not assessed and their respective impacts can thus not be differentiated. The interaction between these two fungal strains thus does not seems to be negative for either strain and even seems positive for *P. citrinum* HEK1. Such an interaction could represent an adaptative strategy to survive under extreme and changing environmental conditions in the same way as dimorphic growth is a common adaptative strategy in fungal extremophiles [70]. The positive impact of this interaction on *P. citrinum* HEK1 is supported by the fact that when spores of *P. citrinum* HEK1 face some stresses, such as cold stresses or ethanol, the yeast cells of *M. guilliermondii* HEK2 grew first and were followed by the subsequent growth of *P. citrinum* HEK1. This may indicate that of *P. citrinum* HEK1 cells benefit from the growth of *M. guilliermondii* HEK2 in challenging environments. Further characterization should be performed to understand how these two organisms interact with each other and how they manage to remain associated overtime. Indeed, the strain *P. citrinum* HEK1 had undergone several cultivation cycles under routine laboratory conditions before the yeast *M. guilliermondii* HEK2 was first observed in particular during the first enrichment phase. Yeast cells must had been associated to the mycelium until the cold stress exposure. The Cryo-SEM images appear to support this, as unrecognized round structures observed alongside the hyphae and spores of *P. citrinum* HEK1 could have been either sexual or asexual cells of *M. guilliermondii* HEK2.

However, despite the co-isolation of these strains from geothermal fluids, their initial origin should be taken with a grain of salt. As much as these strains could have been trapped within the geothermal reservoir for a long period, they could also originate from contamination of the system during the drilling of the wells or during maintenance of the power plant [71], [72]. Nevertheless, these two strains seem to be able to survive under the specific conditions present in the Heemskerk power plant, as they were not isolated directly after the drilling of the wells (2014) [22] or during a maintenance procedure.

Finally, the activity of these strains within the power plant should be assessed, as they can indeed be present either as resistant structures, such as spores, or as active cells. Then, the impact that the presence of these two organisms can have on the geothermal power plant system should be further assessed. It is indeed known that microorganisms can severely impact the functioning of geothermal power plants by inducing mineral precipitation, corrosion of the infrastructures or through the formation of biofilms [15], [16], [17], [18], [73], [74], [75]. However, in geothermal power plants, only the impact of bacteria and archaea has been assessed. The role played by fungi in such infrastructure damages has thus never been investigated, despite fungi being known to participate in pipe clogging, microbially induced corrosion in storage tanks and the biofouling of injectors in the oil and gas industry for instance [76]. As oil and gas industry systems share several similarities with geothermal power plants, such as deep reservoirs, drilling processes or varying temperatures, it can be assumed that fungi may also participate in the microbially induced corrosion or clogging within geothermal power plant systems.

## Acknowledgments

The authors want to thank Chris Boeije, Hester Dijkstra and Laura Wasch for their contribution to the sampling of deep geothermal fluids, as well as Antonio Mucciolo and the Cryo-SEM imaging facility of the University of Lausanne (UniL, CH) for the Cryo-SEM images. Moreover, the authors want to thank Eva Di Francesco, Naïma Mangia and Matys Constantino for their support in the laboratory work.

## Funding

This work was supported by the European Union’s Horizon 2020 research and innovation program within the framework of the REFLECT project under the grant agreement No 850626 and by the Secretariat d’Etat à la formation à la recherche et à l’innovation (SERI) from Switzerland under the contract 22.00148.

## Author contributions

DB: Conceptualization, Data Curation, Formal Analysis, Investigation, Methodology, Data Analysis, Writing – original draft, Writing – review and editing GC: Data Curation, Writing – review and editing WvZ: Sampling, Writing – review and editing SR: Funding acquisition, Supervision, Writing – review and editing SB: Methodology, Writing – review and editing PJ: Conceptualization, Data curation, Formal analysis, Funding acquisition, Investigation, Methodology, Project Administration, Supervision, Writing – original draft, Writing – review and editing

## Data availability

Sanger sequences are available on GenBank under the accession number SUB14533610. Illumina MiSeq sequences used within this study are part of a bigger dataset, available on NCBI as a Sequence Read Archive (SRA) under the accession number SUB13065233. This dataset will be publicly available upon release of the linked publication.

## Conflicts of Interest

The authors declare no conflicts of interest.

## Notes

### Competing Interest Statement

The authors have declared no competing interest.

