## Supplementary Material for "Co-isolation of *Penicillium citrinum* and its cell-switching partner *Meyerozyma guilliermondii* from a geothermal power plant"

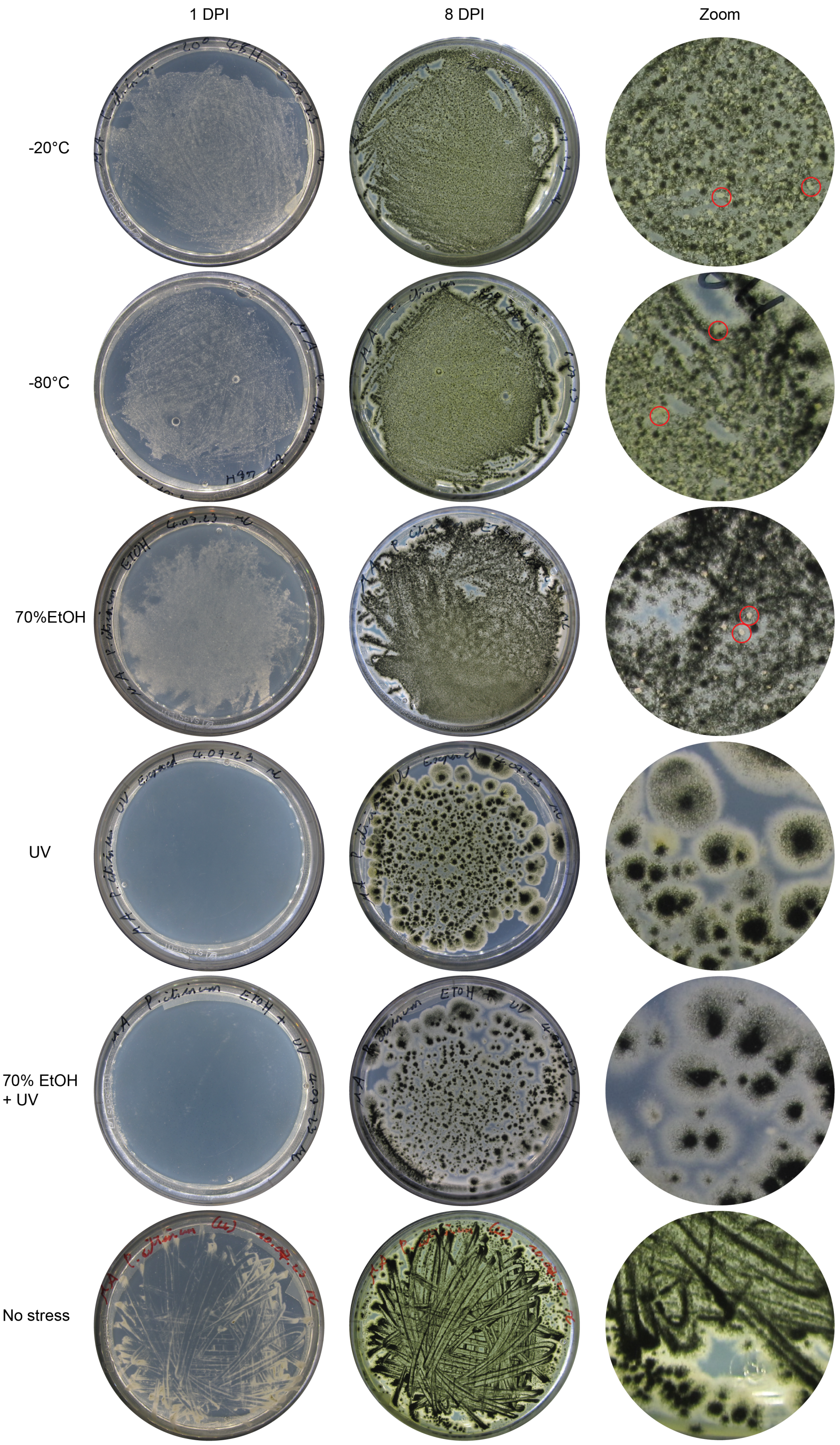


**Supplementary Figure 1 :** Cultures of *P. citrinum* HEK1 under varied stresses at 1 DPI and 8 DPI, except for the "No stress" condition: 1DPI and 7 DPI. Zooms in plates of 7 or 8 DPI are displayed. Yeasts are highlighted by red circles.

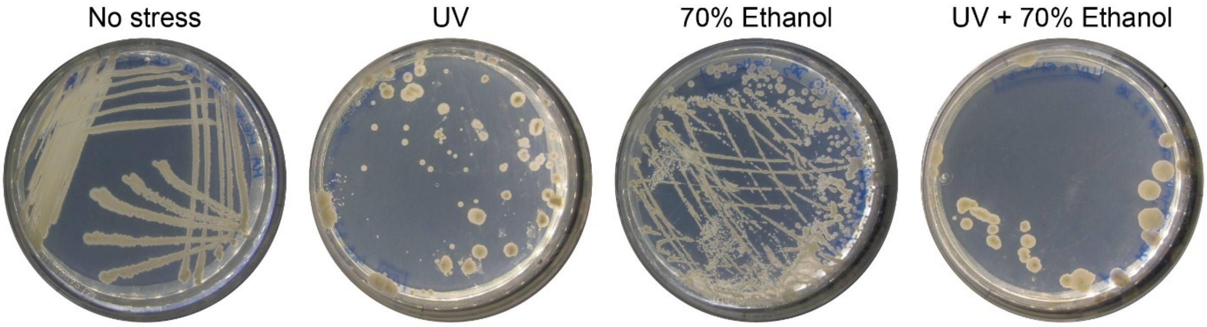


**Supplementary Figure 2 :** Macroscopical aspect of *M. guilliermondii* HEK2 yeast cells under no stress, after UV exposure, after exposure to 70% Ethanol and after a combined exposure to UV and 70% Ethanol.

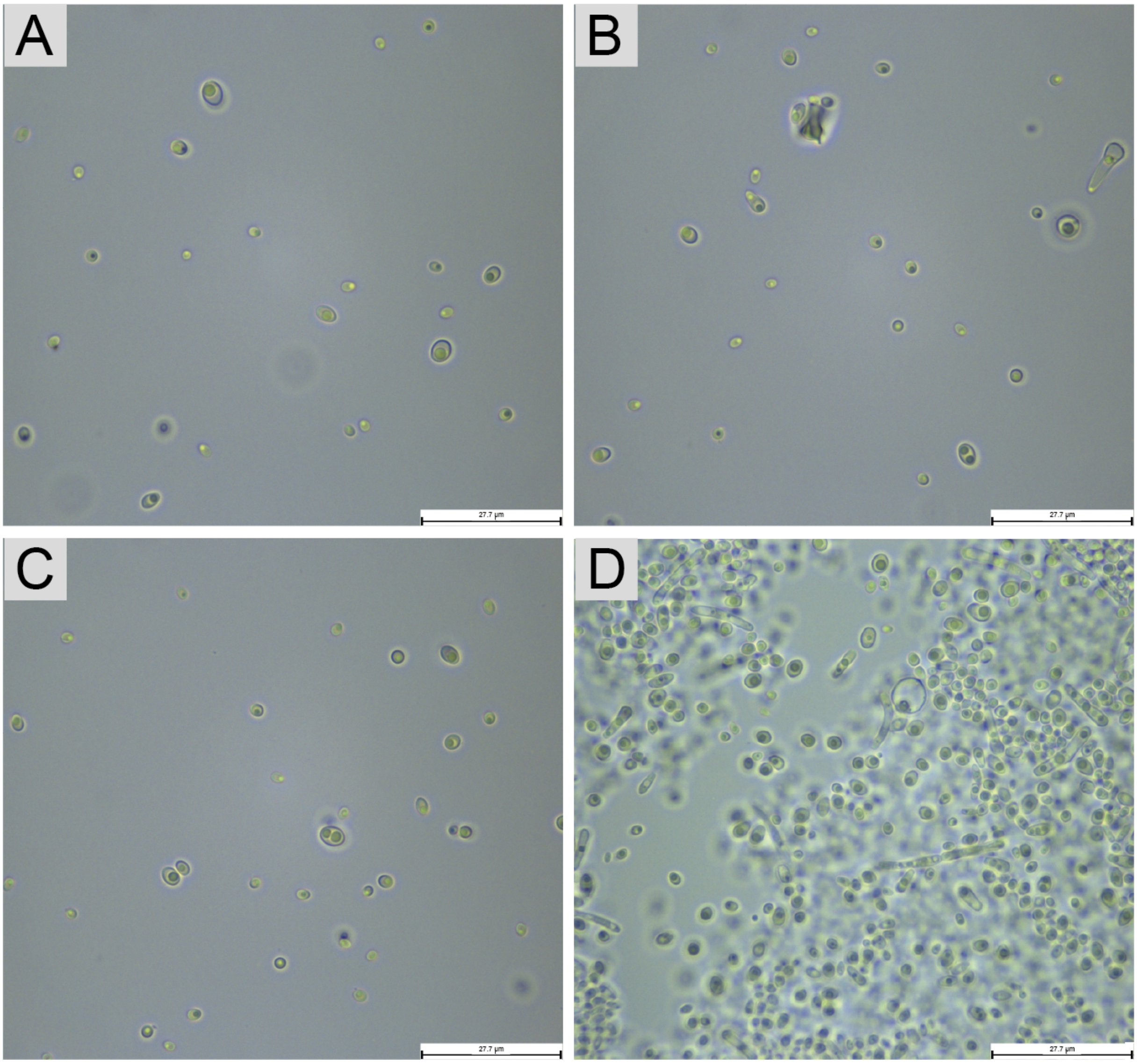


**Supplementary Figure 3 :** Macroscopical aspect of *M. guilliermondii* HEK2 yeast cells A – B) under no stress C) after exposure to 70% Ethanol D) after a combined exposure to UV and 70% Ethanol. Scale bars represent 27.7μm.

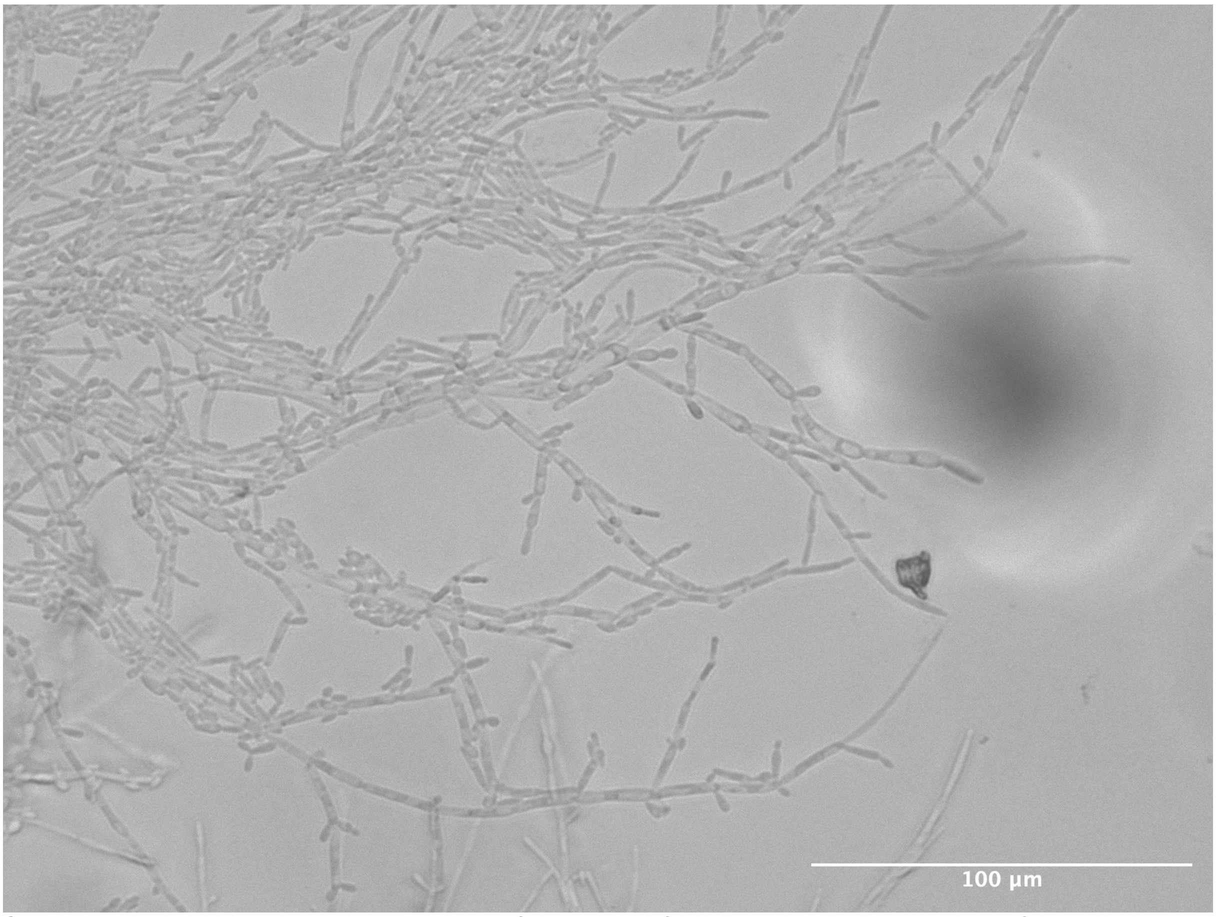


**Supplementary Figure 4:** Time lapse images of the growth of yeast cells into pseudohyphae of *M. guilliermondii* HEK2 on an agar pad at one after 96 h. Scale bar represents 100 µm.

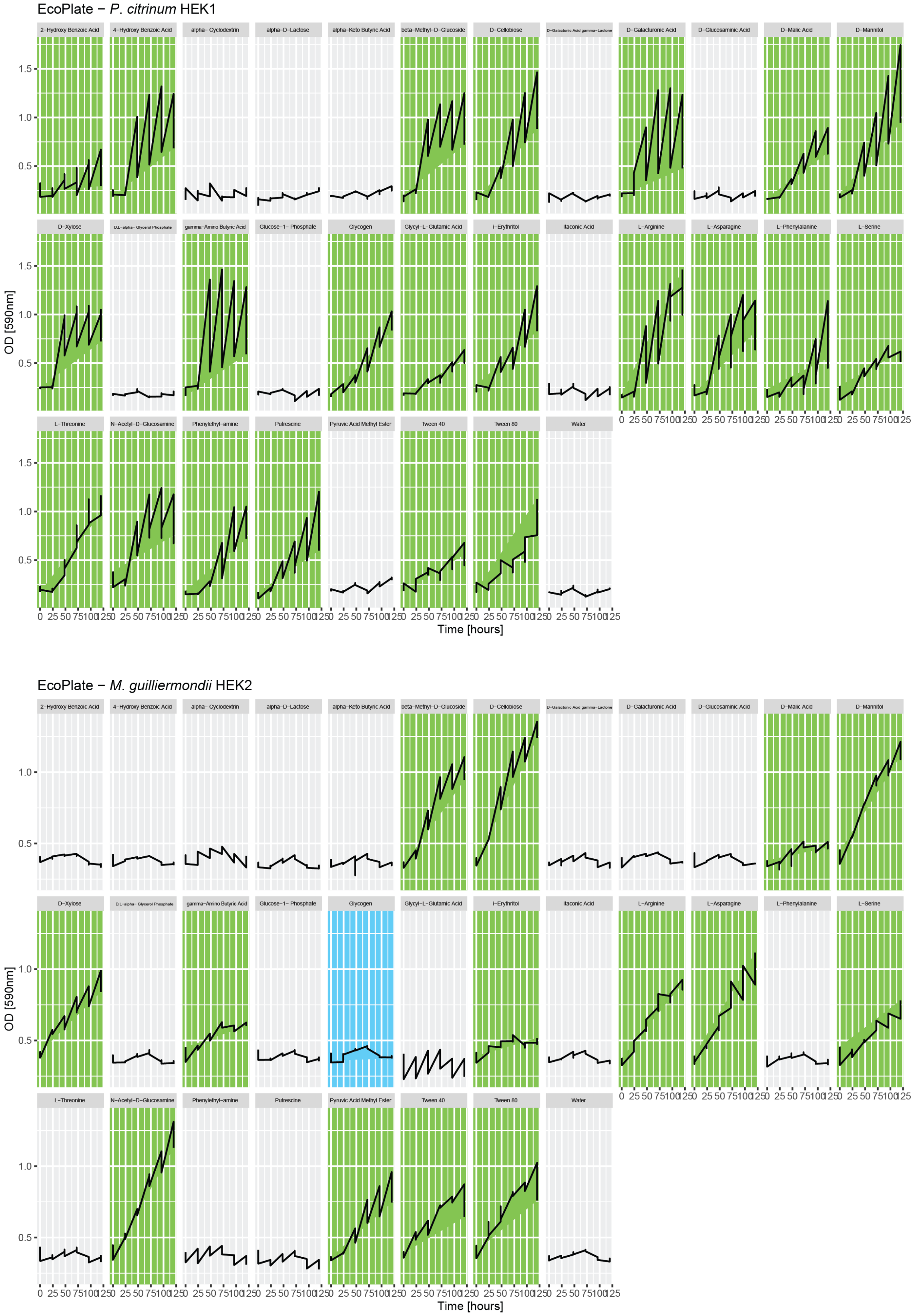


**Supplementary Figure 5:** Growth of the strains *P. citrinum* HEK1 and *M. guilliermondii* HEK2 in different carbon sources in the Biolog EcoPlate™ during 120h. OD measured at 590nm. Time points: 0, 24h, 48h, 72h, 120h. Green: growth confirmed both by the absorbance values and the macroscopic observation of the wells. Blue: growth only observed macroscopically in the wells. The triplicates of the same carbon source were pooled together.

### Media composition

**Supplementary Table 1 :** Composition of the Emerson YpSs Broth, ½ strength medium. Source of original medium: ATCC® Medium 2370 (ATCC®, USA) and [23]. Add components to distilled/ deionized water and bring volume to 1.0 L. Mix thoroughly. Distribute into tubes or flasks. Autoclave for 15 min at 15 psi pressure-121°C.

| | |
|---|---|
| Soluble starch | 7.5 g |
| Yeast extract | 2.0 g |
| K2HPO4 | 0.5 g |
| MgSO4*7H2O | 0.25g |
| pH 7.0 ± 0.2 at 25°C | |

**Supplementary Table 2 :** Composition of the Aspergillus Minimal Medium (AMM). Modified for the H2020 Reflect Project, with an addition of sodium propionate ($C_3H_5NaO_2$), sodium acetate ($C_2H_3NaO_2$) and sodium chloride (NaCl) (LAMUN 2021). Original medium: [77].

| | |
|---|---|
| Glucose | 10g |
| 20x NSS | 50mL |
| 1000x Hutner's trace elements solution | 1mL |
| Technical agar | 18g |
| Sodium propionate | 3 mg |
| Sodium acetate | 76 mg |
| NaCl | 19.45 g |
| MilliQ® $H_2O$ | 1L |
| Adjust pH to 6.5 and autoclave. | |

**Supplementary Table 3:** Composition of the NSS solution: 20x NSS.

| | |
|---|---|
| $NaNO_3$ | 120.0g |
| $KH_2PO_4$ | 30.4 g |
| $MgSO_4 \times 7H_2O$ | 10.4 g |
| KCl | 10.4 g |
| MilliQ® $H_2O$ | 1 L |

**Supplementary Table 4:** Composition of the 1000x Huntner's trace elements solution. For 100mL, dissolve the salts in 80mL of MilliQ® H2O in the order indicated. Most salts will dissolve easily at room temperature, and after addition of Na4EDTA, generally no precipitation is observed. Bring the final volume to 100mL with MilliQ® H2O, autoclave, and store at 4°C. The unadjusted pH will be about 6.5.

| | |
|---|---|
| $ZnSO_4 \times 7H_2O$ | 2.2 g |
| $H_3BO_3$ | 1.1 g |
| $MnCl_2 \times 4H_2O$ | 0.5 g |
| $FeSO_4 \times 7H_2O$ | 0.5 g |
| $CoCl_2 \times 6H_2O$ | 0.16 g |
| $CuSO_4 \times 5H_2O$ | 0.16 g |
| $(NH_4)_6Mo_7O_{24} \times 4H_2O$ | 0.11 g |
| $Na_4EDTA \times 2H_2O$ | 4.94 g |
| MilliQ® $H_2O$ | 100 mL |

**Supplementary Table 5:** Composition of the N-free medium. Original medium: [78].

| | |
|---|---|
| Sucrose | 20 g |
| K2HPO4 | 0.05 g |
| KH2PO4 | 0.15 g |
| CaCl2 | 0.01 g |
| MgSO4. 7H20 | 0.20 g |
| Na2MoO4.2H20 | 0.002 g |
| FeCl3 | 0.01 g |
| CaCO3 | 1 g |
| Agar | 15 g |
| Distillated water | 1 L |
